# CHRONO participates in multi-modal repression of circadian transcriptional complexes

**DOI:** 10.1101/2022.10.04.510902

**Authors:** Priya Crosby, Nicolette F. Goularte, Diksha Sharma, Eefei Chen, Gian Carlo G. Parico, Jon M. Philpott, Rachel Harold, Chelsea L. Gustafson, Carrie L. Partch

## Abstract

The mammalian protein CHRONO is a rhythmically expressed repressor of the circadian transcriptional activator complex CLOCK:BMAL1, and was proposed to be a novel component of the circadian clock. However, lack of specific mechanistic understanding of the activity and function of CHRONO meant that its role within the circadian machinery was opaque. Here we fill this knowledge gap, confirming an evolutionarily conserved minimal repressive domain (MRD) of CHRONO that interacts with specific regions in the BMAL1 C-terminal transactivation domain (TAD) to repress CLOCK:BMAL1 activity. Notably, this binding region overlaps with the binding site for the repressor CRY and coactivators CBP/p300, with CHRONO capable of competing with both of these classical regulators of BMAL1 for TAD binding, highlighting CHRONO as a direct regulator of BMAL1 function.

Additionally, we investigate the interaction between CHRONO and the major circadian repressor, PERIOD2 (PER2). We show that CHRONO reduces PER2 stability through interaction between the CHRONO C-terminus and the Casein Kinase 1 (CK1)-binding domain of PER2. This results in competition between CHRONO and CK1 for binding at this site on PER2, with CHRONO binding inhibiting CK1 phosphorylation of PER2 at the stabilising S662 residue. Taken together, these data show a more substantive and complex role for CHRONO in molecular circadian timekeeping than previously posited, suggesting that CHRONO acts to fine-tune cellular timekeeping by modulating multiple protein-protein interactions that are critical for maintenance of circadian rhythmicity.

## Introduction

Circadian (∼24hr) rhythms allow organisms to synchronise their physiology and behaviour with, and so anticipate, the external day-night cycle. These cell autonomous rhythms regulate a wide range of factors, including sleep/wake cycles, body temperature, metabolism, and hormone levels (Hastings et al., 2003, Reddy and O’Neill, 2010). It has been reported that up to 43% of all protein coding genes are transcribed in a circadian manner in at least one tissue (Zhang et al., 2014, Talamanca et al., 2023). Furthermore, disruption of circadian rhythmicity, as occurs during shift-work and jetlag, is associated with an increased prevalence of a number of disorders, including cardiovascular disease, type II diabetes, some forms of cancer, and compromised immune function (Salgado-Delgado et al., 2013, Scheer et al., 2009, Cuesta et al., 2016). Conversely, disrupted circadian rhythmicity in behaviour is one of the earlier symptoms of several neurodegenerative disorders, including Alzheimer’s and Parkinson’s disease (Li et al., 2017, Saeed and Abbott, 2017). Thus, mechanisms regulating circadian rhythmicity play a significant role in the maintenance of long-term human health.

At the molecular level, the canonical mammalian circadian oscillator consists of a transcription/translation feedback loop, or TTFL. Here, the transcriptional activators CLOCK and BMAL1, which are both basic-helix-loop-helix PER-ARNT-SIM (PAS) domain proteins, form a heterodimeric complex and bind to E-boxes, a DNA response element. This drives expression of many downstream genes, including the transcriptional repressors PERIOD (of which there are three in mammals, PER1/2/3) and CRYPTOCHROME (of which there are two, CRY1/2). PER and CRY dimerise, recruit Casein Kinase 1δ (CK1δ), and translocate to the nucleus, where they form a macromolecular complex with CLOCK:BMAL1 (Aryal et al., 2017), inhibiting their own transcriptional activity through formation of an ‘early’ repressive complex that involves CK1δ-dependent displacement of CLOCK:BMAL1 from DNA (Cao et al., 2021). CK1δ also regulates PER turnover, thereby removing this repressive complex. CRY1 is found with CLOCK:BMAL1 on DNA after the dissolution of this ‘early’ complex, forming the ‘late’ repressive complex; upon CRY1 degradation, CLOCK:BMAL1 is then able to resume its transcriptional activity and the cycle restarts, with this whole process taking approximately 24 hours (Takahashi, 2017, Laothamatas et al., 2023).

CHRONO (also referenced as human C1orf51, GM129 or CIART) has been proposed to be another component of the cellular timekeeping machinery. Previous work identified strong CLOCK:BMAL1 binding within the CHRONO promoter, while high-throughput computational methods simultaneously identified CHRONO expression to be comparable to that of known ‘clock genes’ in a number of parameters (Annayev et al., 2014, Anafi et al., 2014, Goriki et al., 2014, Hatanaka et al., 2010). Furthermore, *chrono* mRNA showed robust circadian oscillations in abundance that are antiphasic to *bmal1* expression and in-phase with *per2* expression (Goriki et al., 2014, Annayev et al., 2014, Hatanaka et al., 2010). Mice with a homozygous deletion of the *chrono* gene exhibited a 25-minute increase in circadian period (Anafi et al., 2014), while *chrono* knockout in U2OS cells resulted in an approximately 2 hour increase in period (Yang et al., 2020), with CHRONO overexpression resulting in period shortening (Goriki et al., 2014). CHRONO also acts as a repressor of CLOCK:BMAL1 activity (Anafi et al., 2014, Annayev et al., 2014, Goriki et al., 2014, Yang et al., 2020). Taken together, CHRONO appears to be a contributing component in the negative arm of the transcription/translation feedback loop, perhaps modulating and refining the activity of the core clock components (Annayev et al., 2014). However, the molecular mechanisms of this modulation are currently unclear, making the specific role of CHRONO within the molecular circadian machinery difficult to define.

Here, we dissect the molecular details of the interaction between CHRONO and its other known core circadian interactors, BMAL1 and PER2, employing biochemical, biophysical, and cellular methods. We confirm a minimal domain of CHRONO required for transcriptional repression via BMAL1, identify critical residues required for interaction of this domain with the BMAL1 transactivation domain (TAD) to carry out its repressive function, and show competition for this binding site with the classical BMAL1 regulators CRY1 and CBP. Additionally, we investigate the interaction between CHRONO and the circadian repressor PER2, showing that the C-terminal portion of CHRONO is sufficient to bind to PER2 within the PER2 Casein Kinase 1 Binding Domain (CK1BD), where it appears to compete with CK1δ. This results in reduced phosphorylation of PER2 by CK1 at the stabilising S662 residue, thereby reducing PER2 stability. Together, this works sheds light on the mechanisms by which CHRONO helps to fine-tune the proteinprotein interactions that are critical to circadian rhythmicity, defining its role in cellular timekeeping.

## Results

### Identification of a minimal repressive domain in CHRONO

In humans, CHRONO is a 385 amino acid protein (**Fig. 1A**) that lacks clearly annotated functional domains. When expressed by transient transfection in HEK 293T cells, the full-length protein is capable of repressing CLOCK:BMAL1 activity at the *per1* promoter in a dose-dependent manner, comparable to the classical CLOCK:BMAL1 repressor CRY1 (**Fig. 1B** (Anafi et al., 2014)). A previous study reported that truncations of CHRONO containing residues 1-284 or 108-385 were sufficient for CLOCK:BMAL1 repression, tentatively identifying a minimal core domain from residues 108-284 (Anafi et al., 2014). We used secondary structure predictions and conservation maps to further investigate these construct boundaries, noting that CHRONO remains highly conserved throughout mammals (**Supplementary Fig. 1A**). However, when other vertebrates, such as fish and birds, are added to the alignment, a conserved core begins to emerge (**Supplementary Fig. 1B**). We found no invertebrate orthologues of CHRONO. Secondary structure predictions found this conserved region to be predominantly structured with numerous alpha helices spanning the length of the construct (**Fig. 1A**). We termed this conserved core of CHRONO (residues 101-245), similar to one recently identified (Yang et al., 2020), the ‘minimal repressive domain’ (MRD).

**Figure 1.**
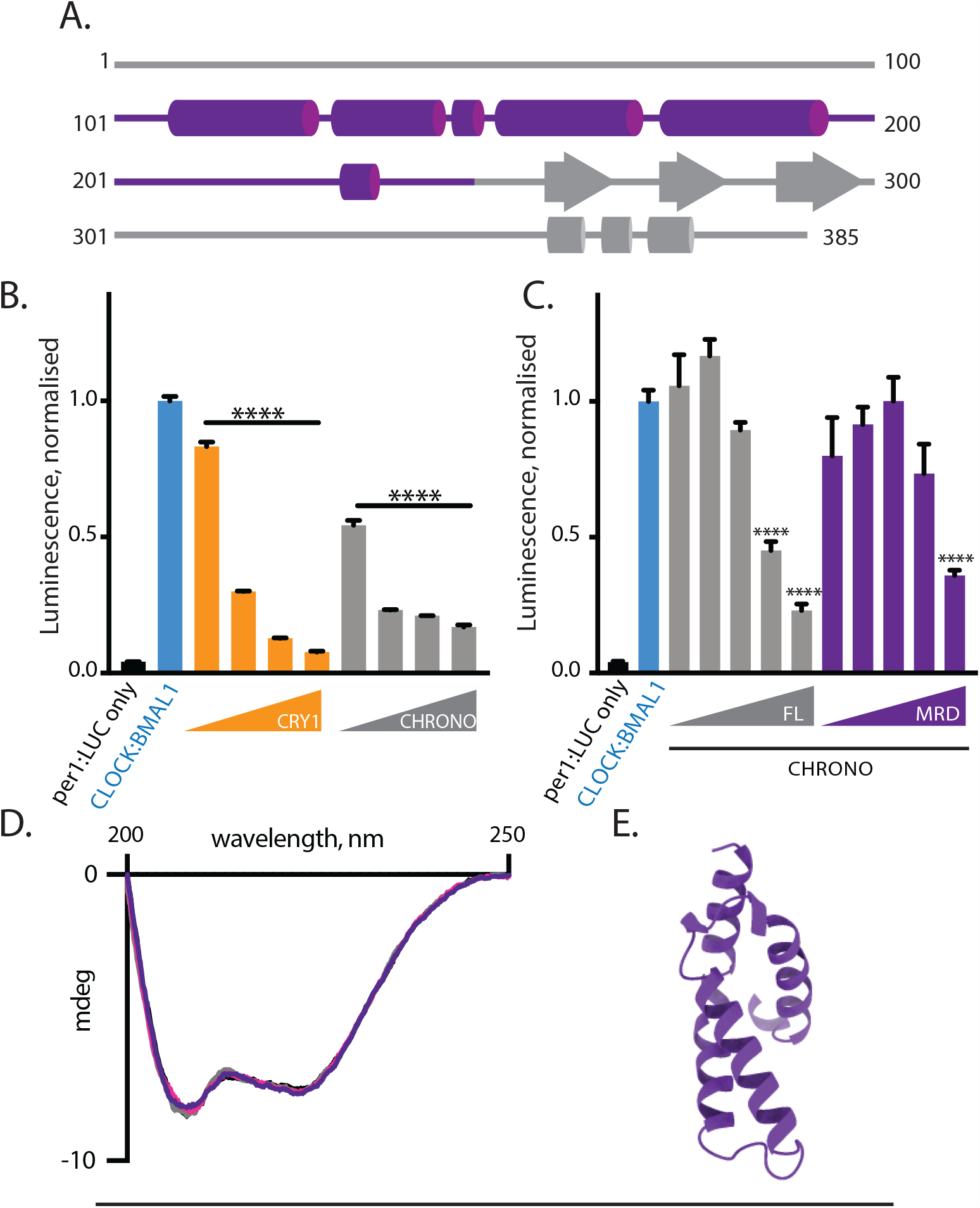
Identification of a minimal repressive domain (MRD) on CHRONO. (A) Schematic of CHRONO domain architecture with secondary structure prediction boundaries from PSIPRED analysis. Minimal repressive domain (MRD) is highlighted in purple. (B) per1:LUC reporter assay of CLOCK:BMAL1 activity in HEK 293T cells transiently transfected with per1:LUC, BMAL1, CLOCK, and increasing concentrations of CRY1 or CHRONO constructs. CRY1 and CHRONO plasmid concentrations range from 0.5-50ng. n=4, mean±SEM. One-way ANOVA with Dunnett’s multiple comparisons test. (C) per1:LUC reporter assay of CLOCK:BMAL1 activity in HEK 293T cells transiently transfected with BMAL1, CLOCK, and increasing concentrations of CHRONO full-length (FL) or CHRONO MRD. Concentrations of CHRONO plasmids range from 0.25-25 ng. n=3, mean±SEM of duplicate measurements from one representative experiment. One-way ANOVA with Dunnett’s multiple comparisons test. (D) Circular dichroism spectrum of CHRONO MRD is indicative of alpha helical secondary structure. 200 μm cell, 4 replicate experiments overlaid. (E) AlphaFold prediction of CHRONO residues 112-196.

Transient transfection of constructs expressing CHRONO full-length (FL) or the MRD, along with per1:LUC, BMAL1 and CLOCK into HEK 293T cells confirmed that the MRD was sufficient to reduce the transcriptional activity of CLOCK:BMAL1, as determined by relative per1:LUC expression (**Fig. 1C**). We do note that the MRD does not repress to the same extent as full-length protein in these luciferase assays, likely attributable to differences in their relative expression levels (**Supplementary Fig. 2A**). As such, all subsequent *in vitro* investigations of CHRONO’s interaction with BMAL1 were performed with the truncated CHRONO MRD construct. We purified the CHRONO MRD recombinantly and confirmed its folding by circular dichroism which, in line with our original structural predictions, indicated alpha helical secondary structure (**Fig. 1D**). This aligns with AlphaFold predictions, which confidently predicts residues 112-196 to form a helical bundle (**Fig. 1E, Supplementary Fig. 2B**)(Jumper et al., 2021, Varadi et al., 2022).

### CHRONO binds to the BMAL1 TAD

The transactivation domain (TAD) of BMAL1, defined from residues 579-626, with a highly conserved core from residues 594-626 (**Fig. 2A, Supplementary Fig. 3A**), binds to both circadian coactivators (CBP/p300) and repressors (CRY1/CRY2) to generate circadian transcriptional oscillations (Xu et al., 2015, Czarna et al., 2013, Takahata et al., 2000). Initial studies reported that CHRONO interacts with a region upstream of the C-terminal TAD, distinct from the CRY1 binding site (Anafi et al., 2014). However, contrary to this, we observed that the purified BMAL1 TAD bound to nickel-bound HisGB1-CHRONO MRD (**Supplementary Fig. 3B**). The affinity of this interaction was then probed using isothermal titration calorimetry (ITC), revealing that the CHRONO MRD and the BMAL1 TAD interact with a K_d_ of approximately 370 nM (369±6 nM) (**Fig. 2B**).

**Figure 2.**
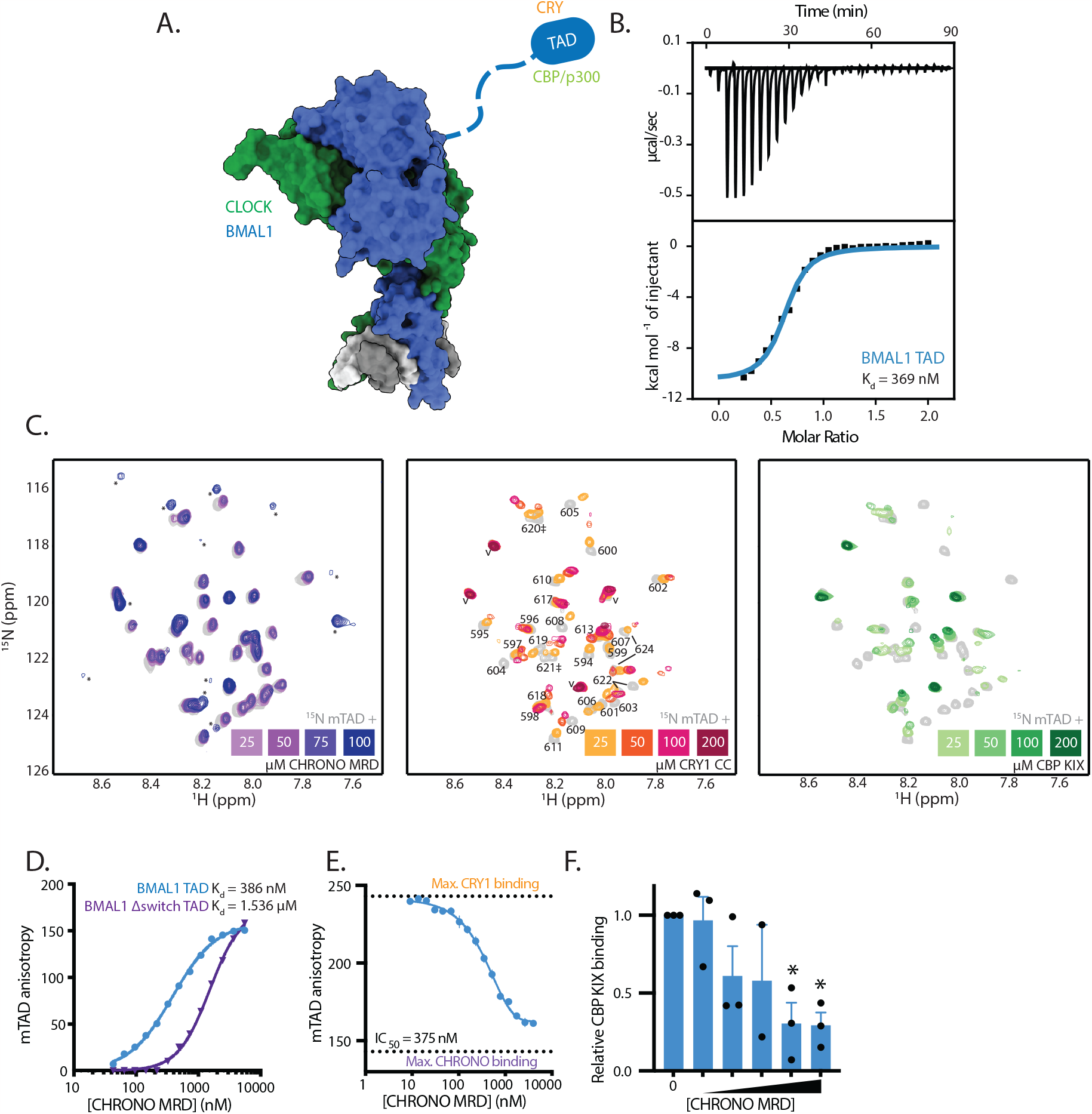
CHRONO binds to the BMAL1 TAD at sites that overlap with coactivator/CRY binding. (A) Model of CLOCK:BMAL1 bHLH PAS-AB bound to DNA, highlighting the BMAL1 TAD as a crucial site for regulation of BMAL1 activity. PDB 4F3L, 4H10. (B) ITC profile for the interaction of His-CHRONO MRD with BMAL1 TAD. Blue line indicates one-site binding model representing best fit to data (K_d_=369±6 nM). (C) ^15^N HSQC spectra showing ^15^N-BMAL1 TAD with titration of CHRONO MRD, CRY1 CC helix and CBP KIX domain. Residues as numbered. ‘*’ indicates new peaks that appear with addition of binding protein. ‘v’ indicates vector residues retained after purification. Residues 620, 621, 622 and 624 all show pairs of peaks due to trans and cis states, indicated by lines or ‡ as appropriate. (D) Fluorescence polarisation with the CHRONO MRD and WT BMAL1 mTAD (K_d_=386±12 nM) or BMAL1 mTAD lacking the switch region (Δswitch) (K_d_=1.536±0.045 μM). n=3, mean±SEM. (E) CHRONO MRD competes with CRY1 PHR for binding at the BMAL1 mTAD with an IC_50_ of 375 nM by fluorescence polarisation. All points performed in duplicate, representative of 3 experiments shown. (F) Density analysis of native gel of TAMRA-BMAL1 mTAD pre-incubated with CBP KIX and increasing concentrations of CHRONO MRD (9.0 nM to 2.3 μM) shows competition for mTAD binding. n=3, one-way ANOVA, Dunnett’s multiple comparisons test. Representative gel shown in Supplementary Fig. 3G.

### CHRONO and CRY1/coactivator binding sites overlap

To pinpoint the CHRONO MRD binding site on the BMAL1 TAD, we collected NMR ^1^H–^15^N heteronuclear single quantum coherence (HSQC) spectra of ^15^N-labeled BMAL1 TAD alone or in the presence of stoichiometric concentrations of the CHRONO MRD (**Supplementary Fig. 3C,D**). We observed differential line broadening upon CHRONO MRD addition, along with a set of new peaks arising from the complex. The dramatic loss in peak intensity was localised throughout the highly conserved core encompassing the alpha helix to the extreme C-terminus of the BMAL1 TAD, including the ‘switch region’, named for *cis/trans* isomerization of the W624-P625 peptide bond (Gustafson et al., 2017) (**Supplementary Fig. 3D,E**). To explore this in more detail, we collected ^1^H–^15^N spectra on this evolutionarily-conserved minimal TAD (mTAD, residues 594-626, **Supplementary Fig. 3F**) in response to titrations of the CHRONO MRD, CRY1 CC helix, or CBP KIX domain (**Fig. 2C**). Chemical shift perturbations and/or broadening are seen at a similar set of mTAD residues with the CHRONO MRD and the other two regulators of this region, demonstrating that they bind similar overlapping regions on the BMAL1 mTAD (**Fig. 2C**). Due to the instability of the isolated MRD at high concentrations, we had to limit its concentration compared to the other proteins. However, consistent with our NMR studies on the larger TAD (**Supplementary Fig. 3C,D,E**), a new set of peaks corresponding to the bound complex began to arise as the MRD and ^15^N mTAD approached equimolar concentrations.

In line with the CHRONO MRD binding the highly conserved core of the BMAL1 TAD, fluorescence polarisation binding assays using a TAMRA-labelled BMAL1 mTAD probe showed that it bound the CHRONO MRD with a comparable affinity to the longer TAD (**Fig. 2D**). This affinity is moderately higher than the affinity of the BMAL1 TAD for the other regulators; for example, full-length CRY1 binds to the BMAL1 TAD with an affinity of ∼1 μM (Czarna et al., 2013) with a similar affinity for the CRY2-TAD complex (Fribourgh et al., 2020), while the CBP KIX domain binds with an affinity of ∼2 μM (Garg et al., 2019, Xu et al., 2015). As with CRY1 and CBP KIX, a construct of the BMAL1 TAD that truncates the switch region led to reduced affinity for the CHRONO MRD (**Fig. 2D**).

The interplay between binding sites for transcriptional co-regulators in this critical region of BMAL1 suggested to us a possible competition-based mechanism, where CHRONO contends with other regulators for TAD binding at different times throughout the circadian cycle. To test this hypothesis, we performed fluorescence polarisation using the TAMRA-labelled BMAL1 mTAD, first making a stable complex with the CRY1 PHR domain, and then titrating in the CHRONO MRD to demonstrate that MRD is indeed capable of directly competing with CRY1 for binding at this region of BMAL1, with an IC_50_ of 375 nM (**Fig. 2E**). Competition for BMAL1 binding has also been previously predicted to occur between CHRONO and the coactivator CBP (Anafi et al., 2014, Yang et al., 2020). Due to similarities in the fluorescence anisotropy values for CHRONO MRD and CBP KIX complexes with the TAMRA-BMAL1 mTAD, we were not able to use fluorescence polarisation to investigate competition between CBP and CHRONO. Instead, we used a native polyacrylamide gel assay, showing that CHRONO MRD could indeed displace pre-bound CBP KIX domain from TAMRA-BMAL1 mTAD (**Fig. 2F, Supplementary Fig. 3G**). Given that CRY1 and CBP KIX also compete for TAD binding (Xu et al., 2015), this demonstrates a multi-way competition that highlights the BMAL1 TAD as a particular hotspot for dynamic regulation of CLOCK:BMAL1 transcriptional activity, with CHRONO a contributing factor in this part of circadian transcriptional activation and repression.

### Specificity of CHRONO for BMAL1 originates from sequence differences in the TAD

BMAL2 is a paralog of BMAL1 that plays a role in mammalian photoperiodism (Wood et al., 2020) but is not capable of maintaining circadian rhythmicity outside of the suprachiasmatic nucleus (Shi et al., 2010, Xu et al., 2015). BMAL2 shares a high degree of sequence and functional conservation with BMAL1 in the structured N-terminal bHLH and PAS domains. However, their disordered C-termini functionally diverge, with the C-terminus of BMAL1 conferring the ability to generate transcriptional oscillations, which BMAL2 cannot (Xu et al., 2015). The minimal TAD of BMAL1 exhibits modest sequence differences from BMAL2, primarily localized to the helical region (**Fig. 3A**). Previous work has shown that CHRONO did not bind, nor was it able to repress, the CLOCK:BMAL2 heterodimer (Anafi et al., 2014). To assess the impact of sequence variation in the BMAL2 TAD on CHRONO binding, we evaluated their interaction by ITC, which showed weak binding between the CHRONO MRD and BMAL2 TAD (**Fig. 3B**). Due to the relatively low heats evolved and the high error value, a dissociation constant could not be calculated.

**Figure 3.**
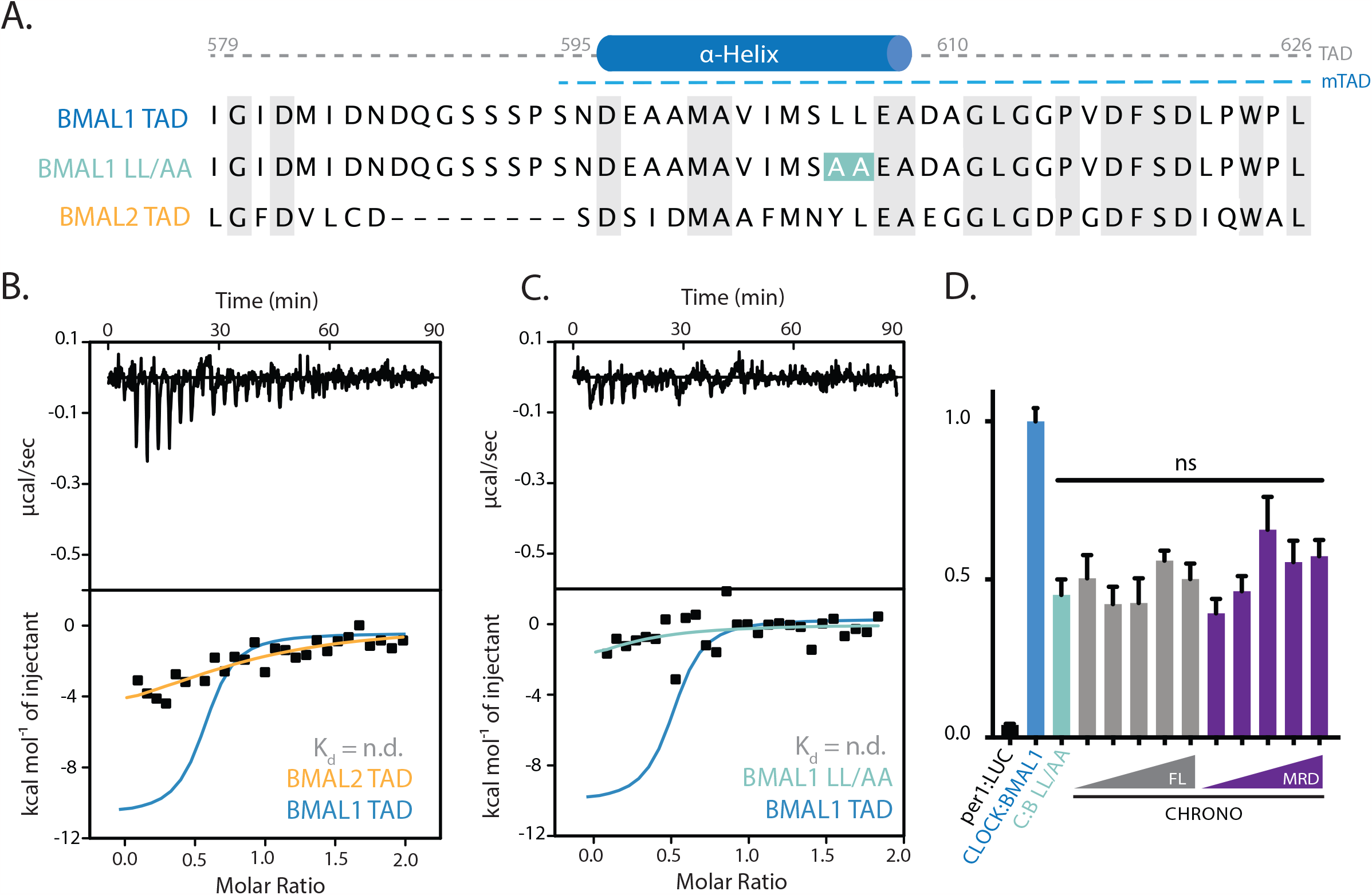
Specificity of CHRONO for BMAL1 originates from sequence differences in the TAD. (A) Alignment of the BMAL1 TAD to its homologous BMAL2 TAD sequence. (B-C) ITC profile of His-CHRONO with BMAL2 TAD and BMAL1 LL/AA shows no substantial interaction. The BMAL1 TAD ITC profile is mapped for comparison (blue). (D) per1:LUC reporter assay of CLOCK:BMAL1 activity in HEK 293T cells transiently transfected with per1:LUC, BMAL1 LL/AA, CLOCK, and increasing concentrations of CHRONO Full-length (FL) or CHRONO MRD constructs. CHRONO plasmid concentrations range from 0.25-25ng/well. n=4, mean±SEM. One-way ANOVA with Dunnett’s multiple comparisons test.

We previously identified a mutant that disrupts binding of transcriptional coregulators to the BMAL1 TAD by introducing L606A and L607A into the TAD alpha helix, referred to as the BMAL1 TAD LL/AA mutant (Xu et al., 2015). This mutant has a dramatically decreased capacity to bind HisGB1-CHRONO MRD by pull-down on nickel resin (**Supplementary Fig. 3H**). ITC with CHRONO MRD and the BMAL1 TAD LL/AA mutant shows that the mutations abrogate binding under these conditions (**Fig. 3C**). Co-transfection of LL/AA mutant BMAL1 into HEK 293T cells with a per1:LUC plasmid confirmed a comparable phenotype in this cellular context: while both full-length CHRONO and CHRONO MRD repressed CLOCK:BMAL1 activity (**Fig. 1C**), the BMAL1 LL/AA mutation eliminated repression by both CHRONO constructs (**Fig. 3D**), as it did for CRY1 (Xu et al., 2015).This confirms that loss of the IxxLL motif in the TAD α-helix, commonly associated with binding of transcriptional coregulators (Dyson and Wright, 2016, Heery et al., 1997), disrupts the binding of CHRONO and the BMAL1 TAD during *in vitro* biochemical assays and thereby also results in a loss of CLOCK:BMAL1 repression by CHRONO in cells. Together these observations confirm the necessity of specific residues within the TAD for CHRONO binding.

### Binding of CHRONO C-terminus regulates PER2 stability

In addition to binding BMAL1, CHRONO has also been reported to interact with the circadian repressor PER2 (Anafi et al., 2014, Annayev et al., 2014, Goriki et al., 2014), although neither the mechanistic basis of this, nor the impact of this on PER2 function, have been investigated. Flag pull-downs using recombinantly-expressed PER2-flag, CK1δ and/or the CHRONO MRD showed that whilst CK1δ bound to PER2 as expected, the CHRONO MRD was insufficient to directly interact (**Fig. 4A**), suggesting that a different region of CHRONO is responsible for binding to PER2. Unlike the full-length protein, the isolated Nor C-termini of CHRONO were expressed at very low levels in HEK 293T cells, but we found evidence that the C-terminus of CHRONO is indeed sufficient to bind PER2 (**Fig. 4B**). Combined with previous work suggesting that the nuclear localisation signal required to traffic CHRONO to the nucleus is found within its N-terminus (Yang et al., 2020), we are now able to begin to annotate multiple functional regions within the CHRONO protein (**Fig. 4C**).

**Figure 4.**
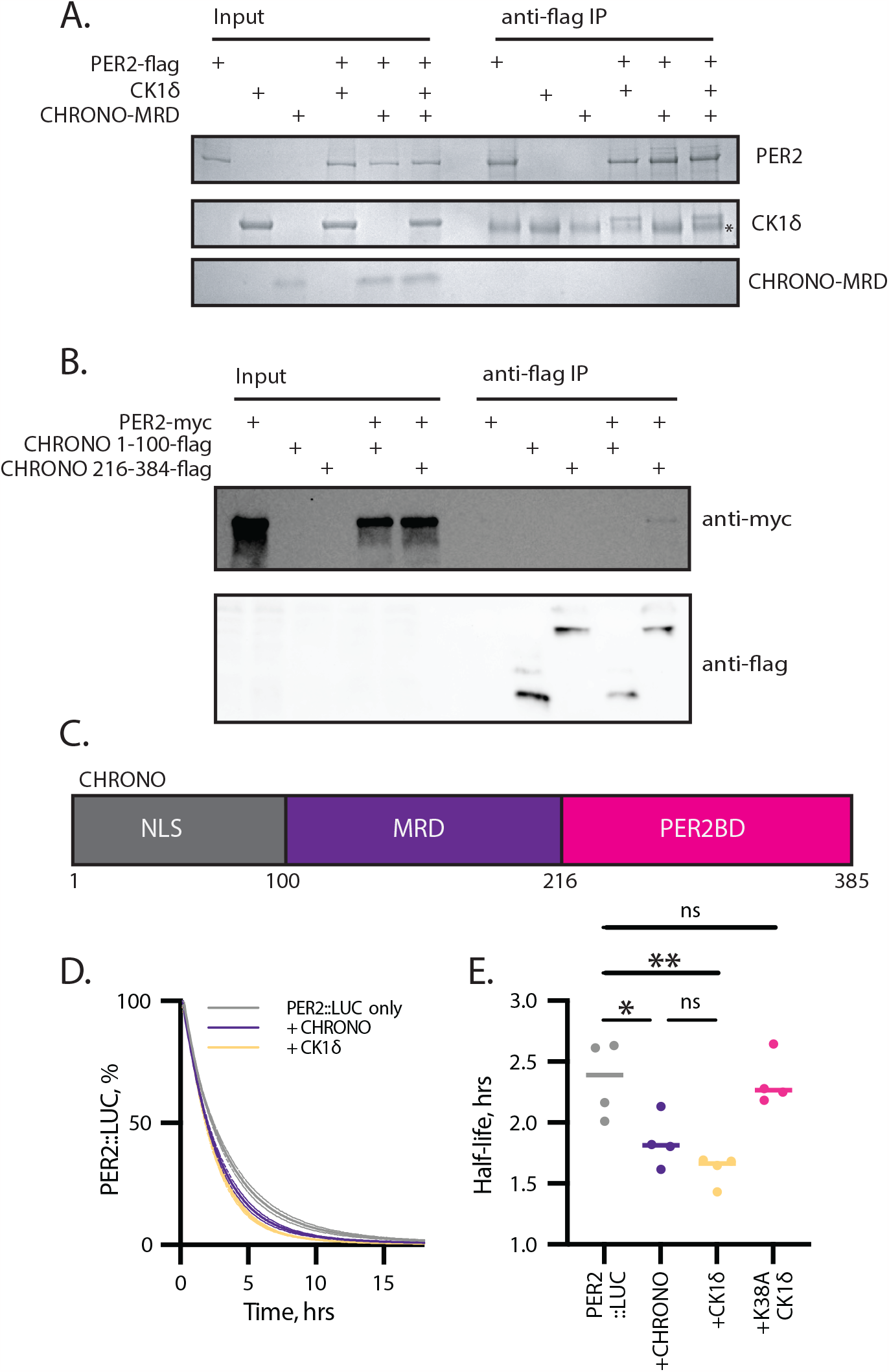
Binding of CHRONO C-terminus influences PER2 stability. (A) Recombinantly expressed full-length PER2-flag pulls down with CK1δ but not the CHRONO MRD. n=2, representative shown. ‘*’ indicates non-specific band. (B) Co-immunoprecipitation of PER2-myc with CHRONO 1-100-flag and CHRONO 216-384-flag shows PER2 to interact with the C-terminal region of CHRONO. n=3, representative shown. (C) Schematic overview of functional regions of CHRONO, highlighting a region containing a nuclear localisation signal (NLS), the minimal repressive domain (MRD) and the C-terminal PER2 Binding Domain (PER2BD). (D) Degradation assay with transient expression of PER2::LUC in HEK 293T cells with CHRONO, CK1δ or catalytically inactive CK1δ K38A shows CHRONO to (E) reduce PER2 half-life following addition of cycloheximide (CHX) at time=0 in a manner comparable to CK1δ. n=4, mean±SEM, representative of 3 independent experiments. Extended Supplementary Fig. 4A.

Classically, the primary post-translational regulator of PER2 stability is Casein Kinase 1 (CK1), particularly the CK1δ isoform (Meng et al., 2010). CK1δ exerts this action on PER2 through stable binding to a dedicated CK1-binding domain (CK1BD) (Eide et al., 2005, Lee et al., 2004). Degradation assays using transient transfection of HEK 293T cells with a PER2::LUC fusion reporter plasmid, along with constructs expressing full-length CHRONO, CK1δ or a catalytically inactive CK1δ K38A mutant showed CHRONO to reduce PER2 half-life in a manner comparable to catalytically active CK1δ (**Fig. 4D,E**, **Supplementary Fig. 4A**). This indicated to us that CHRONO might play an entirely unanticipated role in regulating the repressive arm of the circadian TTFL through regulation of the stability of this essential repressor protein.

### CHRONO competes with CK1 to bind within the CK1-Binding Domain of PER2

Following the surprising observation that CHRONO influences the half-life of PER2, we next sought to determine the region of PER2 to which CHRONO binds. Given CHRONO’s CRY-like repression of BMAL1 through binding to the TAD, we first looked to see if CHRONO also bound PER2 within its CRY Binding Domain (CBD) (Yagita et al., 2002). Cellular co-expression of CHRONO with either full-length PER2 or PER2 truncated just before its C-terminal CRY Binding Domain (PER2ΔCBD) suggests that, unlike CRY, the PER2 CBD is not essential for the interaction between CHRONO and PER2 (**Supplementary Fig. 4B**).

We next made a series of structurally informed truncations in PER2 to see which of these would influence its capacity to bind CHRONO. Truncation of PER2 after the Jα-helix that follows the PAS-B domain (PER2 1-473) abolished CHRONO binding, whilst truncation after the annotated CK1 Binding Domain (CK1BD, PER2 1-763) did not (**Fig. 5A**). More targeted truncations either before the start of the CK1BD (PER2 1-569) or prior to the second conserved motif required for CK1 binding (CK1BD-B) (Eide et al., 2005, Narasimamurthy and Virshup, 2021) also eliminated binding, suggesting that some portion of PER2 from residues 720-763 is required for CHRONO to interact with PER2 (**Fig. 5B,C**). It is worth noting that previous work has shown that CHRONO binds to PER2 but has reduced affinity for the homologues PER1 and PER3 (Annayev et al., 2014, Anafi et al., 2014). Multiple sequence alignment of PER2 residues 720-763 with the comparable regions of PER1 and PER3 (**Fig. 5C**) shows there to be substantial differences in sequence between the proteins in this region, particularly towards the C-terminus, providing a potential basis for this difference in protein-protein interaction.

**Figure 5.**
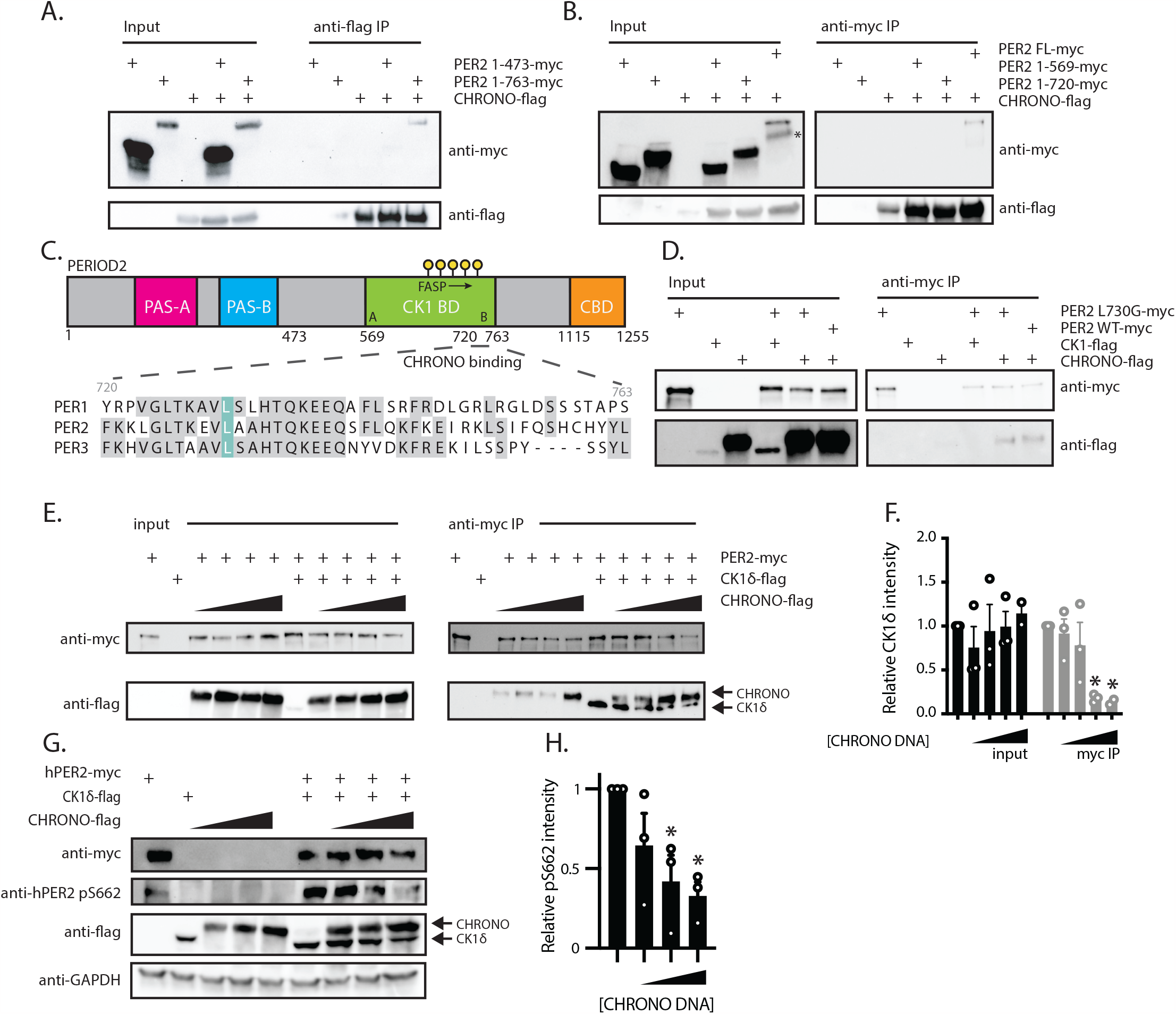
CHRONO binds PER2 within the CK1 binding domain (CK1BD) (A) Co-immunoprecipitation of CHRONO-flag with PER2 truncated after the PAS domains (1-473) or after the CK1BD (1-763). n=3, representative data shown. (B) Co-immunoprecipitation of CHRONO-flag with PER2 truncated before the CK1BD (1-569) or within the CK1BD (1-720). n=3, representative data shown. ‘*’ indicates non-specific band. (C) Schematic of the PER2 domain architecture, indicating the region required for CHRONO binding. ‘A’ and ‘B’ indicate the two CK1BD binding motifs. Inset shows multiple sequence alignment of the three human PERIOD homologues between PER2 residues 720 and 763, with L730 highlighted. (D) Co-immunoprecipitation of CHRO-NO-flag or CK1δ-flag with WT PER2-myc or PER2 L730G-myc. n=3, representative data shown. (E) Co-immunoprecipitation of CK1δ-flag with PER2-myc in the presence of increasing amounts of CHRONO-flag. n=3, representative data shown. (F) Quantification of relative CK1δ intensity, normalised to PER2 intensity. n=3, mean±SEM, one-way ANOVA, Dunnett’s multiple comparisons test. (G) Phosphorylation of hPER2 S662 by CK1δ-flag in the presence of increasing amounts of CHRONO-flag. n=3, representative data shown. (H) Quantification of relative pS662 intensity. n=3, mean±SEM, one-way ANOVA, Dunnett’s multiple comparisons test.

Despite binding within the same region of PER2, the modes of PER2 binding by CK1 and CHRONO must differ, as the PER2 L730G mutation, which abolishes CK1 binding to PER2 (An et al., 2022) has no apparent impact on CHRONO binding to PER2 (**Fig. 5D**). Despite this difference, given the substantial overlap between the identified CK1 and CHRONO binding regions, we posited that these two proteins might influence the other’s interaction with PER2. Transient transfection with constructs expressing PER2, CK1δ and increasing quantities of CHRONO supported this hypothesis, with a reduction in total CK1δ signal observed to co-IP with PER2 as CHRONO levels increased (**Fig. 5E,F**), indicating competition between CK1δ and CHRONO to bind PER2.

Phosphorylation of PER2 by CK1 is a crucial regulator of PER2 stability, with the delicate balance between phosphorylation by CK1 at the stabilising ‘FASP’ region and several destabilising ‘degron’ motifs a critical determinant of PER half-life and overall circadian period (Crosby and Partch, 2020, Philpott et al., 2023). As such, the competition between CHRONO and CK1 for binding to PER2 within the CK1BD suggested to us that CHRONO might act to modulate PER2 stability by influencing PER2 phosphorylation. Western blotting using an antibody raised against the first ‘priming’ phosphoserine in the FASP region (pS662) shows that this in indeed the case, with phosphorylation of this initial serine in the FASP sequence being dramatically reduced in the presence of increasing amounts of CHRONO (**Fig. 5G.H**). According to current understanding of PER phosphorylation, this would result in an increased relative rate of phosphorylation at the various degron sites in PER2, not all of which are CK1-dependent, thereby promoting PER2 turnover (Crosby and Partch, 2020). This is line with our prior observation that CHRONO acts to reduce PER2 stability (**Fig. 4A,B**). This adds a new level to our understanding of circadian regulation by CHRONO, demonstrating that, in addition to directly repressing transcriptional activity through binding to the BMAL1 transactivation domain, CHRONO is also capable of regulating circadian repression indirectly, by altering the stability of the repressor PERIOD2.

## Discussion

Our study builds on previous work that both identified CHRONO and established it to be a component in the molecular circadian oscillator through extensive *in vivo* experiments (Anafi et al., Goriki et al., 2014, Annayev et al., 2014, Hatanaka et al., 2010). CHRONO appears to be part of the negative arm of the circadian TTFL, with its knockout or overexpression influencing circadian period in both animals and cells (Anafi et al., 2014, Goriki et al., 2014, Yang et al., 2020), with its central role attributed to its capacity to repress CLOCK:BMAL1 activity (Anafi et al., 2014, Yang et al., 2020, Annayev et al., 2014). Here we further extend our understanding of CHRONO, probing the molecular details of its interactions that allow it to repress CLOCK:BMAL1, as well as elucidating the nature of its previously unexplored interaction with PER2.

We demonstrate that CHRONO interacts with high affinity at the extreme C-terminus of BMAL1 – the TAD – identifying interactions within the very C-terminal switch region (Gustafson et al., 2017) (**Fig. 2D**) and the α-helix, where conservative mutations abolish binding between the two proteins (**Fig. 3**). Strikingly, we found that this specificity of CHRONO for the TAD of BMAL1 is not extended to the highly related paralog BMAL2. Although overexpression of BMAL2 can partially rescue circadian rhythms of behaviour in whole animals in the absence of BMAL1 (Shi et al., 2010), differences in the regulation of the BMAL1 and BMAL2 TADs shown here for CHRONO, and previously for CRY1 and CBP (Xu et al., 2015), may underlie the persistence of transcriptional rhythmicity at the cellular level that is observed only when BMAL1 is present.

The identification of the BMAL1 TAD as the major site of CHRONO binding to BMAL1, and the competition of CHRONO, CRY and CBP/p300 for this region (**Fig. 2**), adds to the complex role that this region plays in regulating transcription of circadian-controlled genes, with permutations of regulators all vying for the same stretch of amino acids. The BMAL1 TAD likely utilises its intrinsically disordered nature to appeal broadly to circadian regulators, allowing for interactions with both coactivators, CBP/p300, and repressors, CRY and CHRONO, throughout the circadian cycle (Gustafson and Partch, 2015). Regions of a similarly highly dynamic and flexible nature can be found in transactivation domains of important systems other than circadian biology, including ERM/MED25, p53/TAZ2, and SSAP/ZMF1 (Landrieu et al., 2015, Zhang and Childs, 1998, Zhang et al., 1998, Wells et al., 2008). Thus, additional studies of this critical regulatory region will likely provide considerable insight into the dynamic interplay of binding and competition, not only at the BMAL1 TAD, but also systems of transcriptional control.

The nature the interaction between CHRONO and the repressor PERIOD, particularly PER2, discovered here expands CHRONO’s role in circadian repression to a multi-modal one, with CHRONO both directly and indirectly regulating the circadian repressive phase. Classically, CK1 is thought to be the primary regulator of the stability of PER2 (Crosby and Partch, 2020, Eide et al., 2005, Lowrey et al., 2000, Toh et al., 2001), with more recent work from our group and others providing insight into the intricate ‘phosphoswitch’ mechanism by which this works (Philpott et al., 2020, Zhou et al., 2015, Philpott et al., 2023). Briefly, this model considers two CK1 target sites on PER2, a so-called ‘FASP’ region, a series of five serine residues, named for the discovery that mutation of the first of these sites (S662G) results in Familial Advanced Sleep Phase (FASP) syndrome, and a degron, located just after the PAS-B domain at S480 (S478 in mice). Phosphorylation at the FASP increases PER2 stability, whilst the same action at S480 results in recruitment of the E3-ubiquitin ligase β-TrCP and subsequent PER2 degradation, with the balance of phosphorylation between these two regions influencing overall PER2 stability (Zhou et al., 2015). Recent work revealed the mechanism of interplay between these two sites, with the phosphorylated FASP region directly inhibiting kinase activity at the degron, thereby promoting PER stabilisation (Philpott et al., 2023, Philpott et al., 2020). The discovery that CHRONO is able to compete with CK1 for binding to PER2 adds another layer to this model, whereby CHRONO might prevent CK1 from phosphorylating the stabilising FASP region, whilst still allowing for phosphorylation of the degron *in trans*, thereby promoting PER2 degradation (**Fig. 6**). This action would align with previous work showing that CHRONO overexpression shortened circadian period (Goriki et al., 2014) with CHRONO knockout increasing period (Anafi et al., 2014, Yang et al., 2020). It is also possible that CHRONO’s competition with CK1 might also influence the capacity of CK1 to phosphorylate CLOCK to drive dissociation from DNA within the early repressive complex, thereby regulating the dynamics of displacement-type repression that occurs during this circadian phase (Cao et al., 2021).

**Figure 6.**
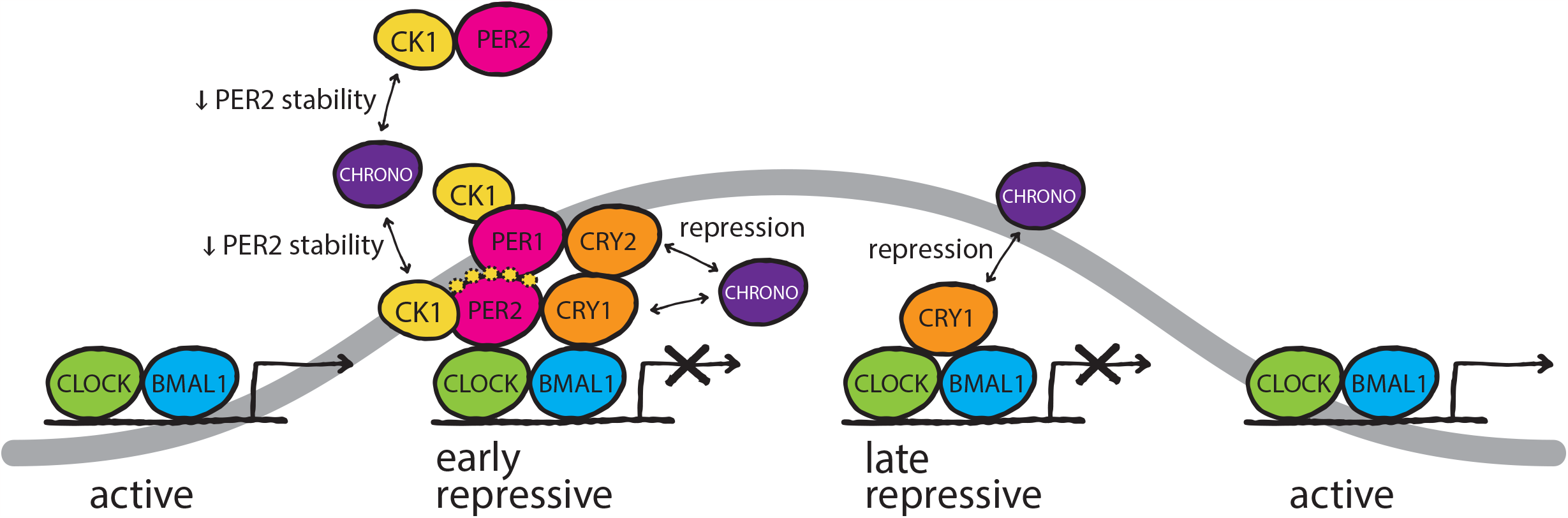
Diagram highlighting the roles of CHRONO in the repressive phases of the mammalian TTFL. CHRONO modulates circadian repression both directly, through competition with CRY1 for binding to the BMAL1 transactivation domain, and indirectly by competing with CK1 for binding to PER2, thereby modulating PER2 phosphorylation and stability. Yellow dashed spots indicate potential phosphorylation events on PER2 that could be modulated by CHRONO.

Overall, this work provides mechanistic insight into the role of CHRONO within the mammalian circadian oscillator, demonstrating its capacity to both directly and indirectly regulate transcriptional repression through both BMAL1 and PER2. This expands our understanding of CHRONO’s role in cellular timekeeping and provides a platform for future work investigating the activity of circadian repressive complexes.

## Materials and Methods

### Multiple sequence alignment and structure prediction

Alignment of protein sequences was performed using COBALT (NCBI) or Clustal Omega using sequences from the NCBI protein database. Alignments were then visualized and annotated using Jalview2 (Waterhouse et al., 2009). Prediction of secondary structure was carried out using PSIPRED 4.0 and AlphaFold.

### Expression and purification of proteins

Human His-CHRONO, HisGB1-CHRONO and HisNusA-CHRONO minimal repression domain (MRD) constructs (aa 101-245) were transformed into the *Escherichia coli* BL21 Rosetta (DE3) strain for recombinant protein expression. Large-scale growths of 1-6L were performed in Luria Broth and induced with 0.5 mM IPTG during logarithmic growth; proteins were expressed for ∼16 hours at 18°C. Cells were resuspended in Buffer A (50 mM Sodium Phosphate pH 7.5, 300 mM NaCl, 10 mM BME, 20 mM Imidazole) and lysed by cell disruption followed by sonication on ice. The extract was clarified by centrifugation at 4°C. The supernatant was filtered through a 0.45-micron membrane, run over a HisTrap affinity chromatography column on a fast performance liquid chromatography (FPLC) instrument, and eluted with a gradient from Buffer A to Buffer B (50 mM Sodium Phosphate pH 7.5, 300 mM NaCl, 10 mM BME, 400 mM Imidazole). His-NusA-CHRONO MRD was then incubated overnight at 4°C with His-TEV to cleave the solubility tag. The sample was subsequently exchanged back into Buffer A and passed over a Nickel-NTA column, eluting into high-salt buffer B (50 mM Sodium Phosphate pH 7.5, 750 mM NaCl, 10 mM BME, 250 mM Imidazole). For all proteins, fractions containing the protein were concentrated and run over a preparative Superdex 75 size exclusion column into CHRONO buffer C (50 mM Sodium Phosphate pH 7.5, 300 mM NaCl, 10 mM BME, 5% glycerol). Purification steps were verified by SDS-PAGE and Coomassie Blue staining to confirm bands at expected molecular weights and assess purity. Protein concentration was assessed by absorbance at 280 nm using a calculated extinction coefficient of 13075 M^-1^ cm^-1^ for the untagged-protein and 24535 M^-1^ cm^-1^ for HisGB1-CHRONO MRD.

Full-length mouse His-PERIOD2-flag (pBP2HF, a kind gift from the Aziz Sancar) was expressed in Sf9 cells. Cells were resuspended in lysis buffer (15 mM HEPES pH 8.0, 250 mM NaCl, 0.1% NP40, 1 mM Na_3_VO_4_, 5 mM NaF, 5% glycerol, EDTA-free protease inhibitors (Roche)) and lysed by sonication on ice. The supernatant was clarified by centrifugation and filtered through a 0.45 μm membrane. The supernatant was then passed over M2 anti-flag affinity gel (Millipore Sigma) under gravity flow, washed with PER2 buffer (1 x TBS pH 7.5, 1mM EDTA, 5% glycerol, EDTA-free protease inhibitors (Roche)) and eluted with 3x flag peptide in PER2 buffer. 1mM TCEP was added to this eluate. The appropriate fractions were then concentrated and run over a Superose 6 Increase 10/300 GL size exclusion column (Cytiva) into PER2 buffer supplemented with 1 mM TCEP to avoid solubly aggregated protein. Purification steps were verified by SDS-PAGE and Coomassie Blue staining to confirm bands at expected molecular weights and assess purity. Protein concentration was assessed by absorbance at 280 nm using a calculated extinction coefficient of 78730 M^-1^ cm^-1^. We found that purified full-length PER2 was prone to forming soluble aggregates upon freeze/thaw or with extended incubation at 4ºC, so all protein in this study was used immediately after purification.

Mouse BMAL1 TAD (residues 579–626), BMAL2 TAD (residues 540–579) and CBP KIX proteins were all expressed as described previously (Xu et al., 2015). The minimal TAD (mTAD) of BMAL1 was cloned from residues 594-626 and purified using the same protocol as the extended TAD.

Human HisGST-CK1δ kinase domain (residues 1-294) preps were grown in the *E. coli* BL21 Rosetta (DE3) strain as described above. Cells were lysed in 50 mM Tris pH 7.5, 300 mM NaCl, 1 mM TCEP, and 5% glycerol using a high-pressure extruder (Avestin) or sonicator on ice. HisGST-CK1δ ΔC fusion proteins were purified using Glutathione Sepharose 4B resin (GE Healthcare) using standard approaches and eluted from the resin using 50 mM Tris pH 7.5, 300 mM NaCl, 1 mM TCEP, 5% glycerol and 25 mM reduced glutathione. His-TEV protease was added to cleave the His-GST tag from CK1δ ΔC at 4°C overnight. Cleaved CK1δ ΔC was further purified away from His-GST and His-TEV using Ni-NTA resin (Qiagen) and subsequent size exclusion chromatography on a HiLoad 16/600 Superdex 75 prep grade column (GE Healthcare) in 50 mM Tris pH 7.5, 200 mM NaCl, 5 mM BME, 1 mM EDTA, and 0.05% Tween 20. Protein concentration was assessed by absorbance at 280 nm using a calculated extinction coefficient of 43320 M^-1^ cm^-1^.

Mouse CRY1 PHR domain (residues 1-491) was expressed in Sf9 suspension insect cells (Expression Systems) as previously described (Parico et al., 2020). Cells were centrifuged at 4 °C at 3,200 × *g*, resuspended in 50 mM Tris, pH 7.5, 300 mM NaCl, 5% (vol/vol) glycerol, and 5 mM BME and lysed in low concentrations of detergent (0.01% Triton X-100), EDTA-free protease inhibitor tablets (Roche), and 1 mM phenylmethylsulfonyl fluoride using a high-pressure extruder (Avestin) followed by brief sonication on ice. After clarifying lysate on a centrifuge at 4°C at 140,500 × *g* for 1 h, protein was captured using Ni-NTA affinity chromatography (Qiagen). Protein was further purified using ion exchange chromatography preceding size exclusion chromatography on a HiLoad 16/600 Superdex 75 prep grade column (GE Healthcare) into CRY buffer (20 mM HEPES pH 7.5, 125 mM NaCl, 5% glycerol, and 2 mM TCEP). CRY1 PHR protein preps were frozen in small aliquots and subjected to only one freeze/thaw cycle.

### Circular dichroism

CD spectra were acquired on a J-1500 CD spectrometer (JASCO). Four independent spectra were acquired with quartz cells of 200 μm pathlength on samples containing 1 mg/mL His-CHRONO in CHRONO buffer (10 mM Sodium Phosphate pH 7.5, 300 mM NaCl, 10 mM BME). Signal from buffer alone was subtracted from the protein spectra.

### Nickel pull-down assays

Nickel pull-down assays were performed by incubating 20 μL Ni-NTA resin, 400 μL Buffer A (50 mM Sodium Phosphate pH 7.5, 300 mM NaCl, 10 mM BME, 20 mM Imidazole) with 2.5 μM HisGB1-CHRONO MRD (bait) and 5 μM of BMAL1 TAD proteins (prey). Samples were incubated overnight at 4 °C with rotation. Resin was pelleted and washed three times, and the bound fraction was eluted with 250 mM Imidazole in Buffer A. Samples were analysed by SDS-PAGE and SYPRO Ruby stain.

### NMR spectroscopy

NMR was conducted at 25°C on a Bruker 800-MHz spectrometer equipped with ^1^H, ^13^C, ^15^N triple resonance, z-axis pulsed field gradient probe. Data were processed using NMRPipe and NMRDraw (Delaglio et al., 1995). Chemical shift assignments were previously made for the BMAL1 TAD using BMRB accession number 25280 (Xu et al., 2015) and manually adjusted for use on the shorter BMAL1 mTAD. ^15^N HSQC titration of 100 μM ^15^N TAD was done by stepwise addition of His-CHRONO MRD in 10 mM MES (pH 6.5), 50 mM NaCl, 10 mM BME. Samples were concentrated to 300 μL final volume and adjusted to a final concentration of 10% (v/v) D_2_O. ^15^N HSQC data were analyzed with NMRViewJ (Johnson, 2004) to extract normalized peak intensities to plot for differential broadening calculations. Differential peak broadening was calculated by dividing the intensity of peaks in a sample of ^15^N TAD containing stoichiometric His-CHRONO by peak intensities of the free ^15^N TAD.

### Isothermal titration calorimetry

Proteins were extensively dialyzed at 4°C in 10 mM Sodium Phosphate pH 7.5, 300 mM NaCl, 10 mM BME using 3 kDa molecular weight cutoff filter dialysis tubing (Spectrum Labs) for 20 hours prior to running ITC. ITC was performed on a MicroCal VP-ITC calorimeter (Malvern Analytical) at 25°C with a stir speed of 155 rpm, reference power of 10 μCal/sec and 10 μL injection sizes. Protein ratios for the cell and syringe for the ITC assays are as follows: 19.2 μM His-CHRONO MRD titrated into 192 μM BMAL1 TAD, 26.7 μM HisCHRONO titrated into 210 μM BMAL2 TAD, and 22.4 μM His-CHRONO MRD titrated into 200.8 μM LL/AA TAD. Data shown are from one representative experiment of two independent ITC experiments were performed for each complex. All data were best fit by a one-site binding model using Origin software with resulting stoichiometry values close to 1.

### Fluorescence anisotropy

The BMAL1 mTAD WT and Δswitch (594-F619Y) probes were purchased from Bio-Synthesis with a 5,6-TAMRA fluorescent probe covalently attached to the N terminus. The C terminus of the Δswitch peptide was amidated, while the WT probe was left as a free carboxyl group to mimic the native C-terminal group of the TAD at L626. Equilibrium binding assays were performed in 50 mM Bis-Tris Propane, pH 7.5, 100 mM NaCl, 0.05% Tween-20, and 2 mM TCEP at room temperature. 20 nM TAMRA-BMAL1 mTAD peptide was preincubated with buffer alone or increasing concentrations of BMAL TAD constructs for 15 min at room temperature before FP analysis. Binding was monitored by changes in fluorescence polarization with a Perkin Elmer EnVision 2103 Multilabel plate reader with excitation at 531 nm and emission at 595 nm. The equilibrium dissociation constant and extent of non-specific binding was calculated by fitting the dosedependent change in millipolarisation level (Δmp) to a one-site specific binding model in GraphPad Prism, with averaged Δmp values from duplicate assays. Data shown are from one representative experiment of three independent assays.

### Endpoint luciferase assays

For per1:LUC reporter assays investigating CHRONO repression, plasmids were transfected in duplicate into HEK293T cells in a 48-well plate using LT-1 transfection reagent (Mirus) with the indicated plasmids: 5 ng/uL Per1:luc, 100 ng each of CLOCK and BMAL1 WT or L606A/L607A, along with the indicated amount of CHRONO full-length (FL) or the Minimal Repressive Domain (MRD); empty pcDNA4 vector was used to normalize total plasmid to 800 ng/well. Cells were harvested 30 hours after transfection using Passive Lysis Buffer (NEB) and luciferase activity determined with Bright-Glo luciferin reagent (Promega). Each assay was repeated three independent times.

### PER2::LUC degradation assays

To investigate the influence of CHRONO on PER2 stability, 200 ng of plasmid expressing the PERIOD2::LUCIFERASE fusion construct under the PGK promoter (pPGK-PER2::LUC, in house) were transiently transfected into HEK293T cells in 35 mm dishes, along with 100ng of either SPORT6-hClorf15-S-tag (CHRONO, a gift from the John Hogenesch lab), pcDNA4B-CK1δ, pcDNA4B-CK1δ K38A, or an empty vector control using PEI at a ratio of 4:1. After ∼16 hours, cells were changed into MOPS-buffered Air Medium (Bicarbonate-free, DMEM, 5 mg/mL glucose, 0.35 mg/mL sodium bicarbonate, 0.02 M MOPS, 2 μg/mL pen/strep, 1% Glutamax, 1 mM luciferin, pH 7.6, 325 mOsm) (Crosby et al., 2017) and moved to a Lumicycle® luminometer (Actimetrics) where bioluminescent activity was recorded at 6 min intervals. After a further 48 hours, 8 μg/mL cycloheximide was added to each dish and returned to recording until the signal reached a baseline.

### Native gels

To demonstrate competition between CBP KIX and CHRONO MRD for the BMAL1 mTAD, native acrylamide gels were poured with 12% acrylamide mix, 0.5x TBE, 1% glycerol, 0.00075% ammonium persulfate and 0.00075% TEMED. Samples were run in 50 mM Tris-HCl, pH 8.0, 100 mM KCl, 1 mM EDTA, 50 ng/ml BSA. TAMRA-mTAD was used at a final concentration of 8 nM, CBP KIX at a final concentration of 105 μM and CHRONO in a four-fold dilution series from 2.3 μM to 9.0 nM. Gels were run at 8 mA and imaged on an Amersham Typhoon™ scanner (Cytiva).

### Co-immunoprecipitation

Plasmids were transfected into HEK293T cells in a 10 cm dishes using PEI 25kDa. Cells were harvested 72 hours later for coimmunoprecipitation with anti-flag M2 affinity gel (Millipore Sigma). Briefly, cells were scraped on ice using 1.5 mL ice-cold PBS and centrifuged for 5 mins at 100 x g at 4°C. The supernatant was removed, and the resulting pellet resuspended in 150 μL ‘lysis buffer’ (20 mM Tris pH 7.5, 150 mM NaCl, 1 mM TCEP, 0.1% NP-40 and EDTA-free protease inhibitors (Roche) and left to lyse on ice for 10 mins. The lysed cells were centrifuged for a further 30 mins at 14000 rpm at 4°C. The resulting supernatant was added to 25 μL pre-washed anti-Flag M2 affinity gel (Millipore Sigma). Tubes were rotated end over end for 2 hrs at 4°C. The gel was subsequently washed three times with 400 μL lysis buffer. Proteins were eluted from resin by addition of 40 μL 2X SDS Laemmli buffer and heated to 65°C for 10 minutes.

### Western blotting

Samples were run on AnyKD™ Mini-PROTEAN TGX™ gels (BioRad) using the manufacturer’s protocol with a Tris-Glycine SDS buffer system. Protein transfer to nitrocellulose for blotting was performed using the Trans-Blot Turbo Transfer system (BioRad), with a standard or high-molecular weight protocol as appropriate. Nitrocellulose was washed briefly, and then blocked for 30 mins at RT in 5% w/w non-fat dried milk (Marvel) in Tris buffered saline/0.05% Tween-20 (TBST). Membranes were then incubated, rocking, with 1:5000 primary antibody (anti-c-myc HRP (Santa Cruz Biotechnology, 9E10, sc-40) or anti-Oct-A HRP to detect flag (Santa Cruz Biotechnology,sc-166355) or anti-pS662 PER2 (abcam, ab206377) diluted in blocking buffer (5% milk, TBST) overnight at 4°C. The following day the membrane was washed for 3 x 10 mins in TBST. For anti-pS662 detection, an anti-rabbit HRP-conjugated secondary antibody (Sigma, A6154) was then added and incubated for 1 hour at RT in blocking buffer, followed by washing for a further 3 x 10 mins in TBST. For all antibodies, chemiluminescence detection was then performed using Immobilon reagent (Millipore), imaged using a ChemiDoc XRS+ imager (Bio-Rad). Quantification was performed using Image Lab Software 6.0 (Bio-Rad).

## Acknowledgements

We thank the Hogenesch lab (Cincinnati Children’s Hospital Medical Center, Cincinnati, OH) and the Sancar Lab (University of North Carolina, School of Medicine) for generously providing the mammalian expression vector SPORT6-hClorf15-S-tag for human CHRONO and the pBP2HF plasmid for full-length mouse PER2 expression respectively. We also thank the members of the Partch and Rubin labs for helpful discussion. This work was funded by NIH grant R01GM107069 and GM1414849 to C.L.P.. G.C.G.P. was funded by a HHMI Gilliam fellowship and the UCSC Graduate Division. P.C. was funded by EMBO ALTF 57-2019.

## Notes

### Competing Interest Statement

The authors have declared no competing interest.

### Summary of Updates

Further experiments to identify the effect of CHRONO on PER2 FASP phosphorylation.

