## Supplementary Figures for "CHRONO participates in multi-modal repression of circadian transcriptional complexes"

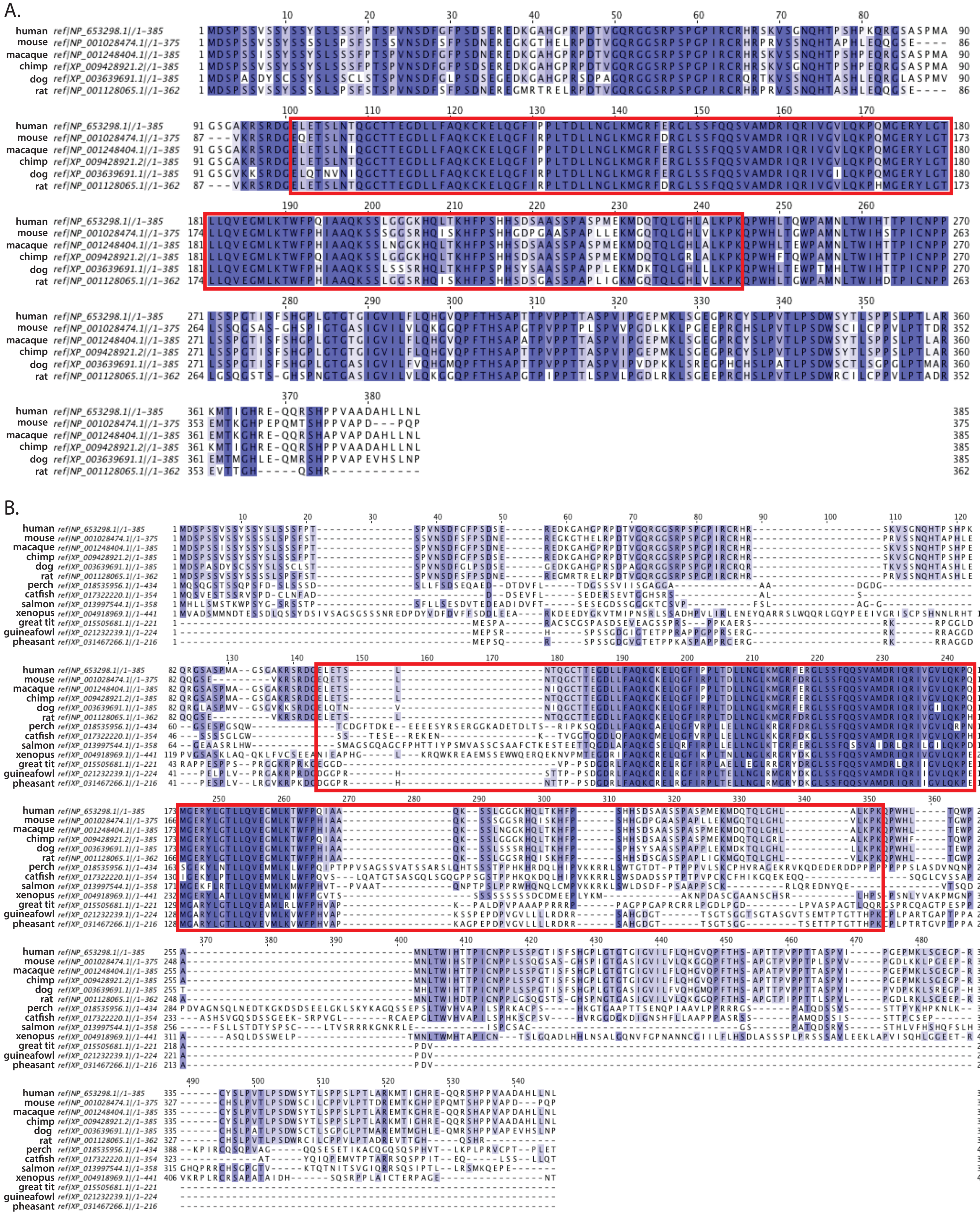

**Supplementary Figure 1. The CHRONO MRD is well-conserved**

(A) Alignment of mammalian CHRONO sequences shows a high level of conservation. (B) Alignment of vertebrate CHRONO protein sequence shows more specific conservation of the core MRD. The defined MRD (residues 101-245 in humans) is highlighted in red.

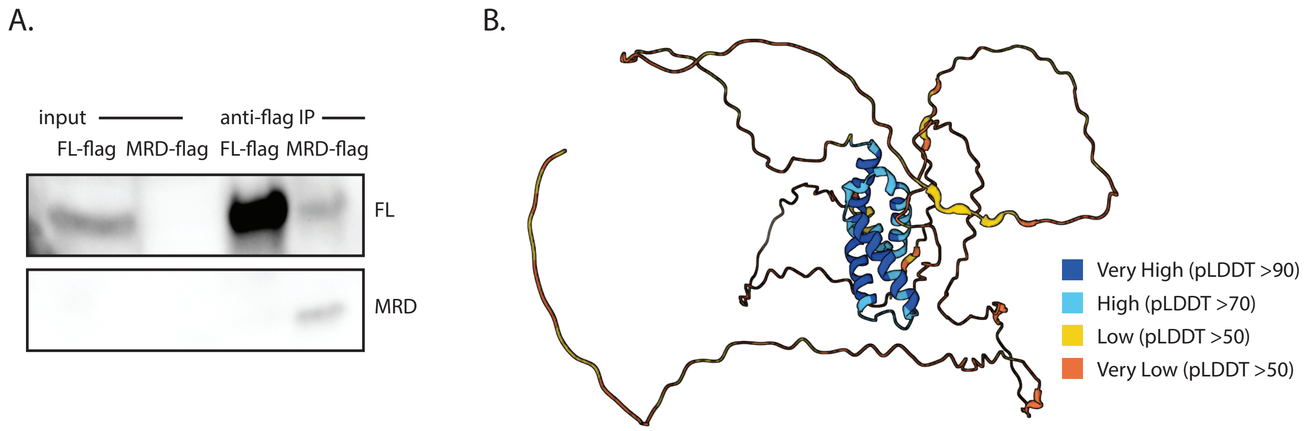


**Supplementary Figure 2. CHRONO MRD predicted structure and relative expression**

(A) Immunoprecipitation of CHRONO MRD-flag and FL CHRONO-flag overexpressed in HEK 293T cells. n=2, representative. (B) AlphaFold predicted structure model of human CHRONO (UniProt Q8N365, AlphaFold AF-Q8N365-F1). Per-residue confidence score indicated.

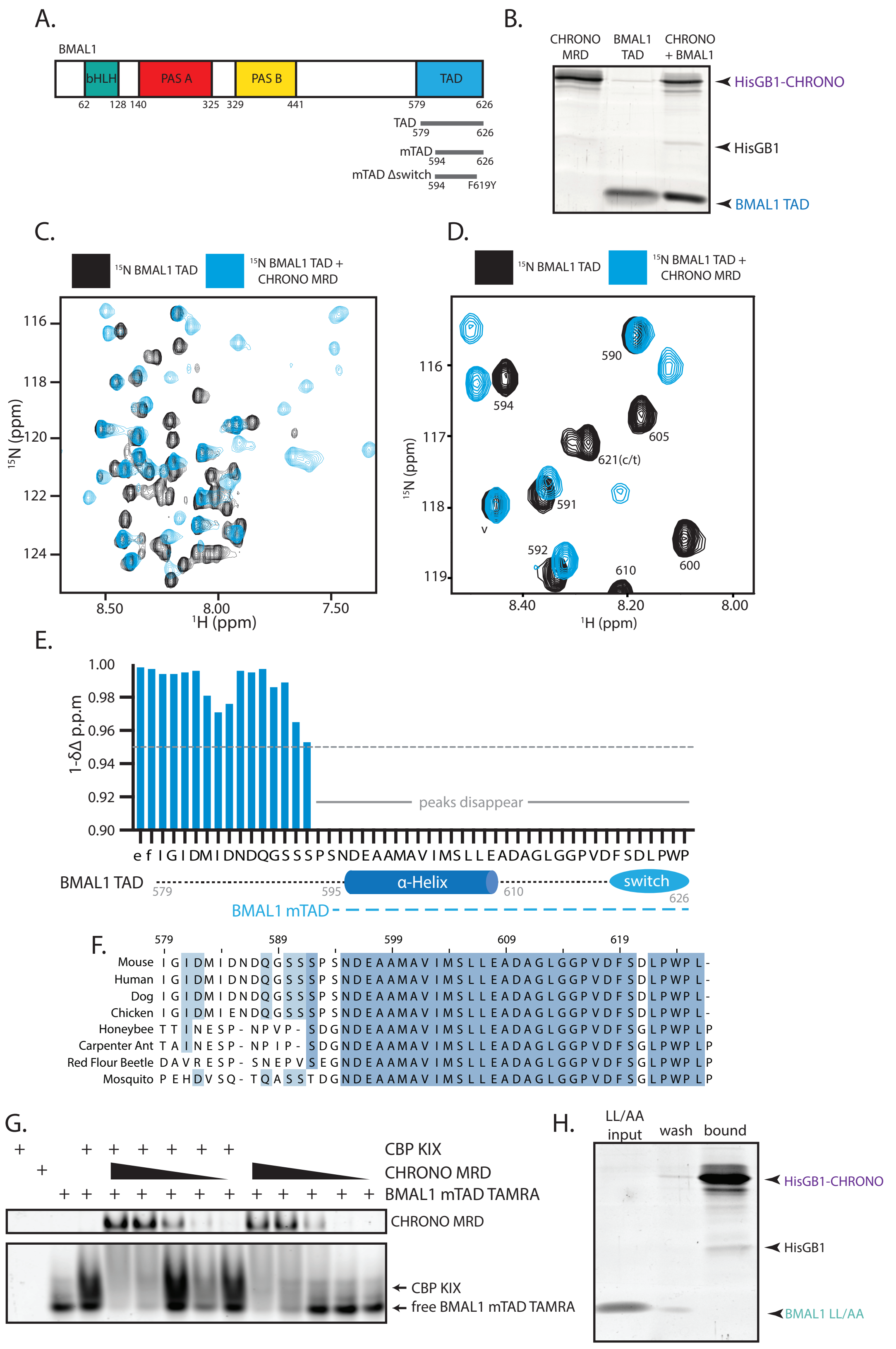


**Supplementary Figure 3. CHRONO binding to the BMAL1 TAD**

(A) Schematic illustrating BMAL1 C-terminal constructs used. (B) In vitro pull-down of purified BMAL1 TAD with HisGB1-CHRONO MRD on Ni-NTA resin. Representative, n=2. (C) $^{15}$N HSQC spectra showing $^{15}$N-BMAL1 TAD with and without CHRONO MRD. (D) Highlighted region shows shifting at some residues, such as 605 (significant shifting) and not at others 590 (minimal shifting). 'v' indicates signal from vector artefact. (E) Differential shift analysis from $^{15}$N HSQC compares peak intensities at 1:1 stoichiometric ratio. A Δintensity close to 1 indicates no binding while a smaller value indicates an interaction or significant change in chemical environment. Proline gives no signal due to the absence of a backbone amide. For other residues, disappearance of peaks indicates change in chemical environment such that a peak comparable to the apo state is no longer detectable. (F) Alignment of vertebrate and invertebrate BMAL1 TAD sequences highlight particular conservation within the C-terminal minimal TAD (mTAD) region. (G) Representative native gel of TAMRA-BMAL1 mTAD pre-incubated with CBP KIX and decreasing concentrations of CHRONO MRD. n=3. (H) *In vitro* pull-down of purified BMAL1 TAD LL/AA mutant with HisGB1-CHRONO MRD on Ni-NTA resin. Representative, n=2.

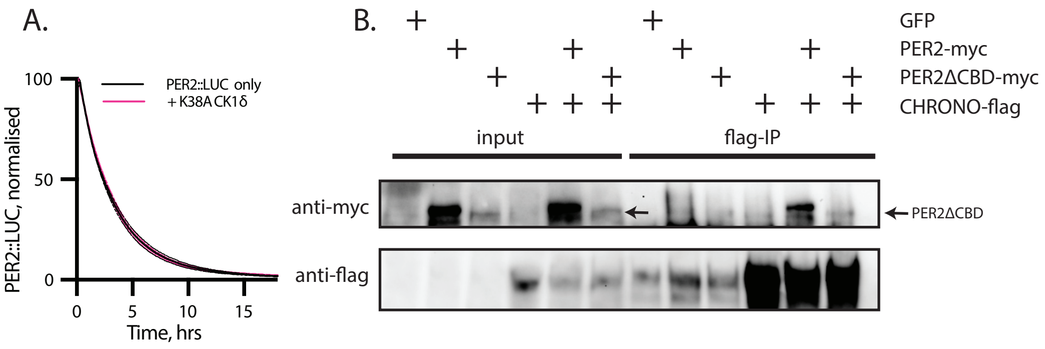


**Supplementary Figure 4. CHRONO binding to PER2**

(A) Degradation assay from co-transfection of catalytically-inactive CK1δ K38A with PER2::LUC. Extended from Figure 4A. (B) Co-immunoprecipitation of CHRONO-flag with full-length PER2-myc or PER2-myc lacking the CRY Binding Domain (PER2ΔCBD) shows that CHRONO is still capable of binding to PER2 in the absence of the CBD. n=3, representative.
